# Endothelial cells release microvesicles that harbor multivesicular bodies and secrete exosomes

**DOI:** 10.1101/2022.10.27.513735

**Authors:** Jennifer D. Petersen, Elena Mekhedov, Sukhbir Kaur, David D. Roberts, Joshua Zimmerberg

## Abstract

Extracellular vesicles (EVs) released by resting endothelial cells support vascular homeostasis. To better understand endothelial cell EV biogenesis, we examined cultured human umbilical vein endothelial cells (HUVECs) prepared by rapid freezing, freeze-substitution, and serial thin section electron microscopy. Thin sections of HUVECs revealed clusters of membrane protrusions on the otherwise smooth cell surface. The protrusions contained membrane-bound organelles, including multivesicular bodies (MVBs), and appeared to be on the verge of pinching off to form microvesicles. Beyond cell peripheries, membrane-bound vesicles with internal MVBs were observed, and serial sections confirmed that they were not connected to cells. These observations are consistent with the notion that these multi-compartmented microvesicles (MCMVs) pinch-off from protrusions. Remarkably, omega figures formed by fusion of MVBs with the MCMV limiting membrane were directly observed, apparently caught in the act of releasing exosomes from the MCMV. In summary, MCMVs are a novel form of EV that bud from membrane protrusions on the HUVEC surface, contain MVBs and release exosomes. These observations suggest that exosomes can be harbored within and released from transiting microvesicles after departure from the parent cell, constituting a new site of exosome biogenesis occurring from endothelial and potentially additional cell types.

## 1 INTRODUCTION

Signaling between cells mediated by secreted membrane-enclosed organelles called extracellular vesicles (EVs) is a widespread form of intercellular communication, evolutionarily conserved from bacteria to plants and animals (Colombo, Raposo, and Thery 2014; Gill, Catchpole, and Forterre 2019). Cells load EVs with a range of bioactive cargos including lipids, membrane proteins, adhesion proteins, cytoskeletal elements, enzymes, signaling molecules, and nucleic acids. Once released into the extracellular milieu, EVs can signal locally or travel long distances in body fluids—such as blood, lymph, cerebrospinal fluid, amniotic fluid—to act on remote tissue targets. Upon reaching recipient cells, specific interactions between EVs and target cells promote binding and uptake by pinocytosis, phagocytosis, endocytosis, or direct fusion with the plasma membrane (van Niel, D’Angelo, and Raposo 2018; Gurung et al. 2021). EV-mediated intercellular signaling is a ubiquitous mechanism occurring under physiological and disease states (Isola and Chen 2017; Cheng and Hill 2022). Plasma EVs are proving sensitive biomarkers of numerous disease states, such as cancer, that can be obtained by relatively non-invasive liquid biopsy (Mitchell et al. 2022). Furthermore, EVs are being developed as vehicles for delivery of therapeutic agents (Sil et al. 2020). Despite these important physiological functions and medical utilities, much remains to be discovered about the biosynthesis of EVs.

It is generally viewed that EVs fall into two categories based on their site of biogenesis. Microvesicles arise at the plasma membrane, by outward budding and pinching off directly from the cell surface. In contrast, exosomes are of endosomal origin, produced when intraluminal vesicles (ILVs) contained in multivesicular bodies (MVBs) are exocytosed upon MVB fusion with the plasma membrane (Mathieu et al. 2019). Microvesicles range in diameter from 50 nm to over 1 μm and there are several types of microvesicles, including ectosomes, microparticles, shedding vesicles, migrasomes, and oncosomes (if released from cancer cell). Apoptotic bodies are a third class of EV, formed when a cell undergoing programmed cell death breaks up into 500 nm – 5 micron-diameter fragments (Xu, Lai, and Hua 2019; Battistelli and Falcieri 2020). In accord with the idea that exosomes represent ILVs that have been released by exocytosis from MVBs, they exhibit a narrower size range between 40-150 nm (Kang, Atukorala, and Mathivanan 2021).

Endothelial cells form the endothelium, a single cell-thick lining of the blood and lymphatic vessels that controls the exchange of oxygen and nutrients between the vessel contents and underlying tissues (Ricard et al. 2021). In their role on the “front lines” of vessels, exposed to circulating body fluids, endothelial cells release a significant proportion of the EVs found in blood (Mathiesen et al. 2021). EVs released by the resting endothelium contribute to its role in supporting vascular homeostasis which includes maintenance of the antithrombogenic surface of the vessels (blood fluidity) and vasodilation, inhibition of inflammation, cell survival and angiogenesis (Trisko et al. 2022).

To gain a better understanding of the mechanisms and structural aspects of basal release of EVs from resting endothelial cells, we used thin section electron microscopy (EM) to examine (HUVECs) and look for structural features consistent with microvesicle budding from the plasma membrane and exocytic release of exosomes from MVBs. Cells were preserved by ultra-fast freezing, which is optimal for capturing fast events like exocytosis, and processed by a freeze substitution protocol optimized for plasma membrane enhancement (Walther and Ziegler 2002).

In thin sections, groups of protrusions were observed on the otherwise smooth HUVEC plasma membrane that were often branched and contained vesicular organelles, including MVBs with ILVs. Beyond cell peripheries, vesicles that contained MVBs were observed, suggesting that they were microvesicles that had pinched off from the protrusions, diffused, and occasionally adhered to the coverslip. Serial sections through the presumptive microvesicles on the coverslip confirmed that they were not connected to cells by cellular extensions and that they contained MVBs with ILVs. Further examination revealed omega figures, the structural hallmarks of exocytosis, occurring between MVB-like vesicles inside the microvesicles and their limiting membrane. On occasion, such omega figures contained small vesicles that were identical to ILVs. These observations support the notion that microvesicles containing multiple membrane compartments (referred to as multi-compartmented microvesicles, or MCMVs) pinch off from MVB-containing protrusions at specialized sites on the cell surface. MCMVs contain MVBs and that apparently can release exosomes after transiting away from the parent cell.

## 2 METHODS and MATERIALS

### 2.1 HUVEC culture on Aclar coverslips

Primary umbilical vein endothelial cells (HUVECs) were obtained from American Type Culture Collection (ATCC, Manassas, VA, USA; #PSC-100-013). HUVECs between passages 4-6 were used in this study, grown to ∼70% confluence, and screened for mycoplasma contamination (ATCC, Kit #30-1012K). Cells were plated on Aclar 33C plastic film (Electron Microscopy Sciences (EMS), Hatfield, PA, USA) which separates easily from polymerized resin (compared to glass) without damaging the bottom of the cells. To plate HUVECs on Aclar, the plastic sheet of Aclar was cut into about 12 mm-wide rectangular coverslips to fit into wells of 12-well plate using clean scissors. After cutting into desired size and shape, coverslips were washed at least ten times with distilled water and then at least ten times with 70% ethanol. Coverslips were then placed into wells of a 12-well plate and rinsed at least ten times with cell culture grade sterile water (ThermoFisher Scientific, Waltham, MA, USA), and then once with endothelial cell culture medium (ScienCell, Carlsbad, CA, USA, Cat. No. 1001) which contains 5% fetal bovine serum, 1% endothelial cell growth supplement, and 5% Penicillin/Streptomycin antibiotic solution. Cells were plated (30,000 cells per well) and maintained for 2-3 days in an incubator at 37°C and 5% CO2. Medium was changed every 48 hours.

### 2.2 Plunge freezing of cultured HUVECs

To preserve cells for EM, cultured HUVECs on Aclar coverslips were plunge frozen by hand in liquid ethane. Ethane was liquified in the “ethane pot” of a Vitrobot Mark IV plunge freezer (ThermoFisher Scientific) following manufacturer’s instructions. A vessel of liquid nitrogen was placed next to the ethane pot for transfer of frozen coverslips to storage containers under liquid nitrogen. To freeze, a coverslip was gently withdrawn from well of 12-well plate with pair of clean, fine-tipped forceps (Dumont #5, EMS). Holding the coverslip vertically, excess culture medium was wicked away from the edges of the coverslip using Whatman #1 filter paper (EMS). Total blotting time did not exceed 5 seconds, with careful observation to avoid drying. After blotting, the coverslip was inverted over the liquid ethane, and plunged into the liquid ethane by hand as quickly as possible. The coverslip was immediately transferred to liquid nitrogen and stored under liquid nitrogen until processing by freeze substitution.

### 2.3 Freeze-Substitution of cultured HUVECs on Aclar coverslips

The freeze substitution (FS) process used was a slightly modified version of a protocol developed to enhance preservation and staining of lipid membranes (Walther and Ziegler 2002). FS staining cocktail was prepared in 20 mL glass scintillation vials with plug caps (EMS) and contained: 0.2% osmium tetroxide, 0.1% uranyl acetate (UA) in glass distilled acetone with 5% water. To prepare the FS cocktail, working under a fume hood, 12 mg of UA (EMS) was added per vial, followed by 600 μl of 4% aqueous osmium (EMS) and vortexed for 30 seconds to dissolve. Then, 11.4 mL of acetone from a freshly opened bottle of glass distilled acetone (EMS) was added to each vial. The vial was tightly capped, and contents mixed well by swirling. Then caps were loosened and placed in a liquid nitrogen bath until the FS cocktail was frozen solid. Liquid nitrogen was poured directly in the vials, and then frozen coverslips bearing cells were transferred into vials while being maintained under liquid nitrogen. One coverslip was processed per vial. Vials were loosely capped and transferred to the chamber of the freeze substitution machine that contained about ½ inch-deep bath of ethanol to improve thermal conductivity and was pre-cooled to -112°C (EM AFSII, Leica Microsystems, Wetzlar, Germany). Within about 10 minutes in the freeze substitution machine, the liquid nitrogen inside the vials was evaporated and vial caps were tightened. The following FS program was applied: warm from -112°C to -90°C, then hold at -90°C for 8 hours, then warm to 20°C for 36 hours.

### 2.4 Room temperature processing and embedding of freeze substituted HUVECs

When the FS program was completed, vials were removed from machine and processed at room temperature. Coverslips were rinsed with acetone three times and then stained with 0.5% tannic acid in acetone for 45 minutes, protected from light. Tannic acid was removed by four acetone rinses using freshly opened bottle of glass distilled acetone. Cells were infiltrated with a graded series of EmBED812 resin (EMS) in acetone (50%, 75%, 95%, and 100% resin twice.) Each infiltration step lasted for at least one hour. The coverslips were embedded as follows: A square “backing piece” of Aclar was cut into a square and placed in the top or bottom half of a 35 mm plastic Petri dish. Then, the coverslip to be embedded was removed from the scintillation vial with clean forceps, and excess resin was allowed to drain from the coverslip for a few seconds. Then the coverslip was placed cell-side-up on the Aclar backing piece in the Petri dish. A gelatin capsule (EMS, size 0) was filled with freshly prepared 100% resin until it was level with the rim of the open capsule, and any bubbles were allowed to float to the surface and removed. Then the uncapped, resin-containing gelatin capsule was inverted and placed directly on top of the cell-side of the coverslip. Two or three gelatin capsules were placed side-by-side to cover most of the surface of the coverslip, and the entire assembly was put into the 60°C oven to for 48 hours to polymerize.

### 2.5 Serial thin sectioning of embedded HUVECs

After polymerization, the backing piece of Aclar was removed and the gelatin capsules attached to coverslips were separated from each other with a jewelers saw into individual blocks. Often the gelatin capsule could be sawed into 3-4 pieces to allow sectioning of multiple areas per block. After trimming away excess resin with a double-edged razor blade, the Aclar was removed by inserting the edge of the razor blade at a corner, between the Aclar and the resin surface, and lifting carefully to flick away the Aclar coverslip, revealing the smooth bottom surface of embedded cells. The block face was then trimmed for sectioning. Sections were cut using an ultramicrotome (EM UC7, Leica Microsystems) to a thickness of 70 nm using a diamond knife (DiATOME, Hatfield, PA, USA). Typically, four to five serial sections were picked up per 2 × 1 mm formvar and carbon coated slot grid (EMS). Because the Aclar coverslip, and therefore the resin surface, was not perfectly flat, the knife often grazed the portions of the block face and missed others at first, with successive sections becoming more complete until the knife fully entered the resin. The partial sections were collected and used to determine the distance from the coverslip of serially collected thin sections. Sections on the grids were post-stained with 3% Reynolds lead citrate (EMS) for 5 minutes.

### 2.6 Transmission EM of serial thin sections and image processing

Thin sections were viewed using a Tecnai T20 transmission electron microscope (ThermoFisher Scientific) operated at 200 KeV. When a structure of interest was discovered that spanned multiple sections, the structure was tracked back in the serial sections to the first section in which it appeared, and then imaged in consecutive sections moving up in the Z-axis away from the coverslip. Images were collected with an NanoSprint1200 side-mount CMOS detector (Advanced Microscopy Techniques, Woburn, MA, USA). Adobe Photoshop 2022 (San Jose, CA, USA) was used to adjust images for display by applying a Gaussian blur of 0.5 pixel-radius, followed by adjustment of grey levels, brightness, and contrast. Serial section images were aligned using the TrakEM2 plugin of Fiji image analysis software (Schindelin et al. 2012).

## 3 RESULTS

### 3.1 HUVECs display a localized cluster of plasma membrane protrusions

To look for sites of EV biogenesis occurring from resting HUVECs using thin section electron microscopy (EM), cells were cultured on Aclar plastic coverslips and then preserved by ultrafast freezing in liquid ethane. Frozen cells were processed by freeze substitution (Walther and Ziegler 2002), and embedded in epoxy resin. The Aclar coverslip was separated from the hardened resin, leaving the cells cleanly transferred from the coverslip to the resin, with the surface of the resin corresponding to the bottom of the cells. Seventy nanometer thick sections were cut in the *en face* orientation to cells, parallel to where the coverslip had been. As the diamond knife entered the resin, all sections were collected, including partial sections that had only grazed portions of the resin surface. Sections were collected in order so that structures spanning multiple sections could be interpreted, and distance from the surface of the coverslip in the z-axis could be determined.

Cells were prepared at ∼70% confluence, allowing isolated peripheries of individual cells where EVs might be emerging to be clearly examined. Thin sections revealed the characteristic elongated HUVEC morphology (Figure 1a). Cells displayed mostly smooth plasma membrane surfaces, however, occasionally a site spanning several micrometers of the cell surface membrane became irregularly contoured (Figure 1a, boxed area shown in 1b). Evaluation at high magnification revealed the area to be composed of a cluster of protrusions emanating from the surface of the HUVEC.

**FIGURE 1:**
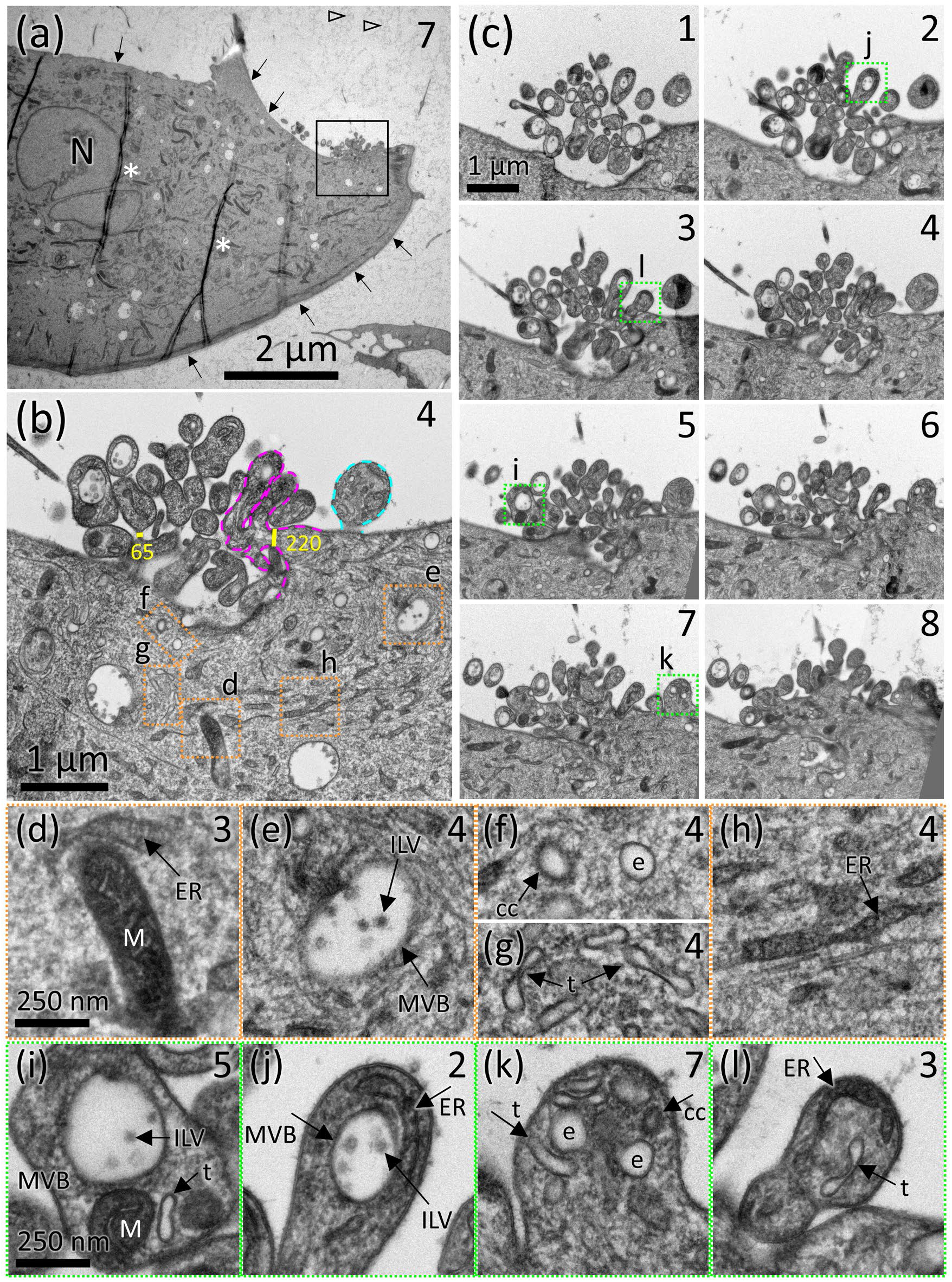
HUVECs display a localized site of membrane protrusions that contain MVBs and other membrane-bound organelles. (a) A low-mag transmission EM view of a portion of a HUVEC in a 70 nm-thick section cut in the *en face* orientation, showing its elongated cell morphology and predominantly smooth plasma membrane (arrows). The number in the upper right corner indicates the number of the section in the series of serial sections shown in (c), N indicates the nucleus. Additional serial sections through this area are shown in Supplemental movie 1. Asterisks are placed to the right of two wrinkles in the plastic section; open arrowheads point to fuzzy material that is typically observed in freeze-substituted culture medium. (b) Boxed area in (a) from serial section 4 shows the cluster of membrane protrusions. A magenta dashed line outlines a branched protrusion; turquoise dashed line outlines an unbranched protrusion. Yellow lines indicate width in nm across protrusion necks. (c) Eight consecutive serial sections through the protrusion site with the serial section number indicted in the upper right (the greater the number, the higher above the coverslip in the Z-axis). (d-h) Enlarged views of cytoplasmic organelles from orange boxed areas in (b) that can be identified based on their ultrastructure include: MVBs containing ILVs, mitochondria (M), endoplasmic reticulum (ER), round endosome (e), clathrin coated vesicle (cc), and tubular endosome (t). (i-l) Enlarged views of individual protrusions from green boxed areas in (c). Organelles identical to those observed in the cytoplasm are indicated.

To explore the three-dimensional organization of this ‘protrusion site’, images of the area were taken across consecutive serial sections and aligned (Supplemental movie 1). The first eight serial sections starting from the surface of the coverslip (Figure 1c) show that the site consists of numerous protrusions, many of which were branched (Figure 1b, magenta outline). Thin necks connecting protrusions to the cell are visible in serial section 4, about 280 nm above the coverslip surface. In sections that occur below or above the level at which the protrusions connect to the cell, the protrusions cut in cross-section appeared to be a cloud of vesicles, hovering next to the cell. For example, Figure 1c, section 1 shows the cross-section of tips of protrusions hanging down towards the coverslip, below where they connect to the cell. At protrusion branch points and where protrusions connected to the cell, diameters often constricted to 65 - 200 nm wide necks (Figure 1b, yellow bars). Some protrusions spanned only 3-4 serial sections (280 nm in z-height with respect to the coverslip), while others extended higher above the surface of the coverslip for about 10 sections (700 nm in z-height). Additional serial sections through this protrusions site are shown in Supplemental movie 1. Two more examples of protrusion sites on other HUVECs are shown in Supplemental figures 1 and 2 and corresponding Supplemental movies 2 and 3. These examples show protrusions that span over 1200 nm (17 sections) in height. Interestingly, multiple protrusion sites were not observed on a single cell, suggesting there may only be one per cell. Based on the frequency that protrusion sites were observed by thin section, which is sampling only a small portion of the HUVEC surface, it is estimated that most cells have a protrusion site.

### 3.2 Organelle-rich protrusions contain MVBs with ILVs, endosomes, ER, and mitochondria

Organelles in the cell cytoplasm could be identified based on their stereotypical EM ultrastructure (orange boxed areas in Figure 1b, enlarged in Figure 1 d-h). In the protrusions, membrane-bound organelles of matching structure were observed, including MVBs containing ILVs, endosomes (round, tubular, and clathrin coated), and endoplasmic reticulum (ER). Occasionally, mitochondria, recognizable by their distinct cristae, double membranes and dark electron dense staining were also observed in the protrusions ((green boxed areas in Figure 1c, enlarged in Figure 1 i-l).

### 3.3 Vesicles were present on the coverslip surface that were not connected to cells and contained MVBs and other membrane-bound compartments

Based on the protrusion morphology—thin necks connecting protrusions to the cell and at branch points—it was hypothesized that the protrusion site is a specialized site for the assembly and budding of EVs. Because the membrane would be derived from the HUVEC plasma membrane, such an EV would qualify as a microvesicle. Furthermore, should a protrusion containing membrane-bound organelles pinch off and diffuse into the extracellular milieu, it would create an microvesicle containing multiple compartments, i.e., a multi-compartmented microvesicle (MCMV). If so, diffusing MCMVs may occasionally contact and stick to the coverslip, and remain attached during preparation for EM. To explore this possibility, areas beyond cell peripheries on seemingly empty expanses of coverslip were searched in the *en face* sections that were closest to the coverslip surface (illustrated in Supplemental figure 3 and corresponding Supplemental movie 4).

In such areas, presumptive MCMVs were found beyond the periphery of cells. An example shown in Figure 2a was located about 7 μm from the nearest cell. A complete set of 10 serial sections through this MCMV was obtained starting at from the very first section that contained the MCMV within the first 70 nm of the surface of the coverslip, to the 10^th^ and last section in which the MCMV appeared (Figure 2b and Supplemental movie 5). Examination of the serial sections confirmed that the MCMV was not connected to a neighboring cell by a membrane extension, nor were remnants of detached membrane connections observed.

**FIGURE 2:**
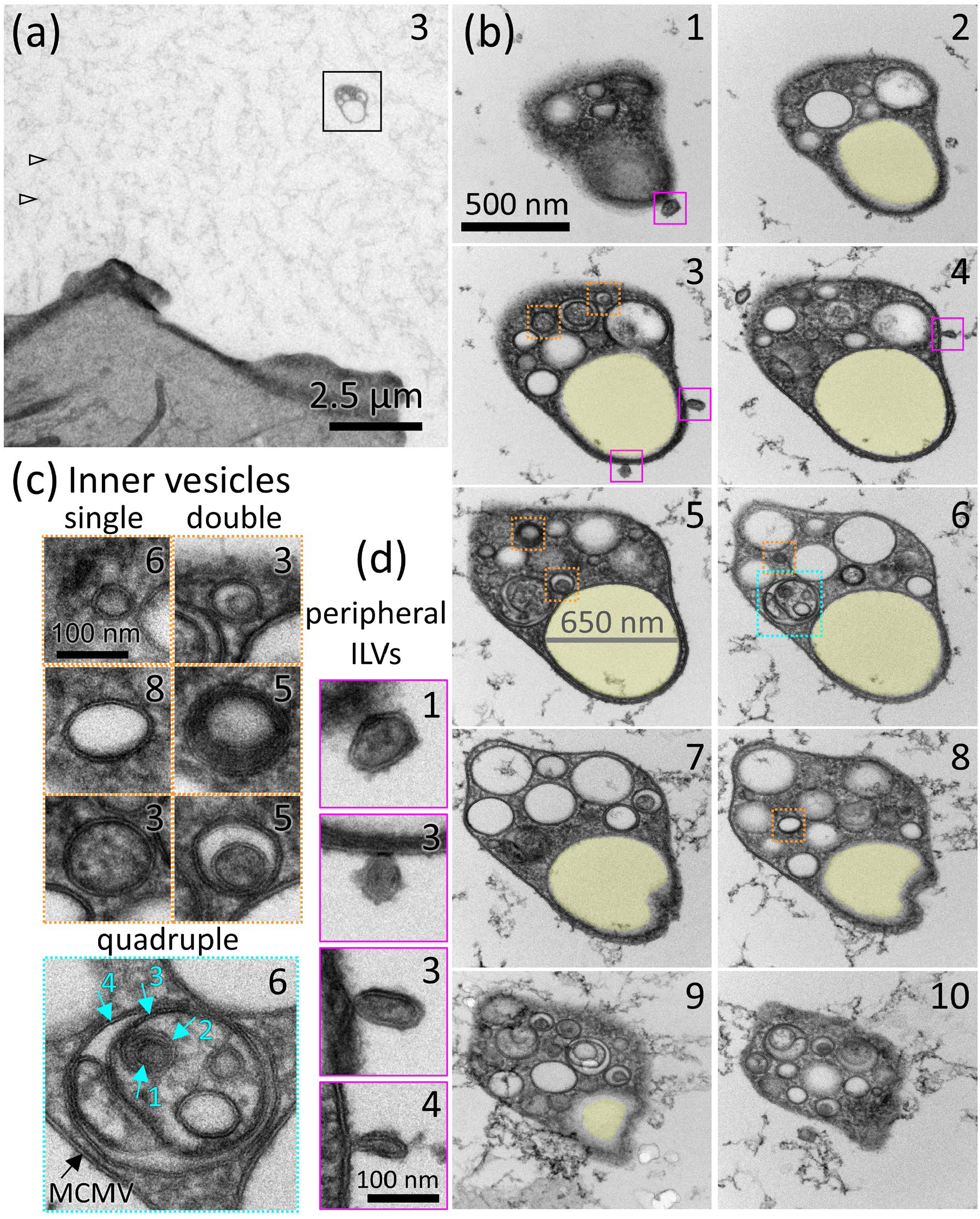
MCMVs are found on the coverslip surface and are completely independent from neighboring cells. (a) A low-magnification view of a presumptive MCMV (boxed) located 7 μm beyond the periphery of the nearest cell. Arrows point to fuzzy material that is typically observed in freeze-substituted culture medium. The complete series of serial sections through the MCMV boxed in (a) is shown in (b) and Supplemental movie 5. The number in the upper right corner indicates the number of the serial section (ascending number indicates moving higher in the Z-axis with respect to the surface of the coverslip). ILV-like vesicles stuck to the outer periphery of the MCMV are boxed in magenta and shown enlarged in (d). Examples of vesicles inside the MCMV are boxed in orange or turquoise and shown in (c). A 650-nm diameter MVB-like vesicle that spans almost the entire series of sections but is devoid of ILVs is shaded yellow in (b). (c) Examples of MCMV inner vesicles that are single vesicles or double vesicles, boxed orange, or a quadruple vesicle, boxed in turquoise. Turquoise arrows indicate 4 layers of membrane of the quadruple vesicle, inside the MCMV limiting membrane (black arrow), amounting to 5 membrane layers. Inner vesicles displayed various combinations of dark, clear, or granular lumens. (d) ILV-like vesicles dangling from the periphery of the MCMV (magenta boxes in (b))

### 3.4 MCMVs contain MVBs with ILVs and other membrane-bound compartments

Serial sections through MCMVs demonstrated that they contained round and tubular membrane-bound compartments in addition to a dense cytoplasm (Figure 2b). The vesicular compartments varied in diameter from 45 to 650 nm. Lumens of the vesicles below 200 nm in diameter varied from electron lucent to electron dense with a smooth or granular texture (Figure 2c). The larger compartments were most often clear, like the lumens of MVBs (Figure 1 b, e, i, j). Often, an MCMV inner vesicle contained another vesicle, i.e., a vesicle inside a vesicle, or double vesicle (Figure 2c). The lumens of the double vesicles often differed in texture or electron density from the vesicle in which it was enclosed (see Figure 2c, double vesicles from section 5). On occasion, MCMV inner vesicles contained multiple vesicles. A quadruple vesicle in Figure 2c from section 6 shows an inner vesicle that contains a round 250 nm vesicle and a 60 nm thick tubule, both with granular textured lumens. The 250 nm round vesicle contains 4 additional vesicles with diameters ranging from 50 - 75 nm with smooth light to dark grey lumens. One of these vesicles contains a sixth, 45 nm vesicle, also with a smooth grey lumen. In this configuration, there are 5 layers of membrane between the contents of the sixth vesicle and the extra-MCMV milieu.

Also present in the MCMV shown in Figure 2b, is a 650 nm sub-compartment (shaded yellow), along with some ∼300 nm compartments that are similar in roundness and clear lumens to MVBs, but they do not contain any ILVs. The presence of putative empty MVBs inside MCMVs raised the possibility that MVBs might be capable of releasing their ILVs (to become exosomes) from an MCMV after it has transited away from its parent cell. Intriguingly, vesicles identical to ILVs were sometimes observed stuck to the outside surface of an MCMV, such as those boxed in magenta in Figure 2b (enlarged in Figure 2d). This observation provided circumstantial evidence for ILV secretion (exosome release) from MCMVs.

### 3.5 MVBs in MCMVs release exosomes by exocytosis of ILVs

If exosomes are released from MCMVs, MVBs containing ILVs must be observed inside MCMVs, and evidence of membrane fusion between the MVB the MCMV membrane must be detected. Sections of MCMVs were examined, and examples of MVBs, indistinguishable from MVBs observed in cell cytoplasm were observed (Figure 3c, and d section 8, upper c’ and d’). The MVB in Figure 3c appears to be undergoing an outward budding event, suggesting active remodeling of MVBs inside MCMVs (inset Figure 3 upper c’). Other MVB-like vesicles inside MCMVs were observed (Figure 3a, orange boxes, and upper 3a’, and Supplemental Figure 4).

**FIGURE 3:**
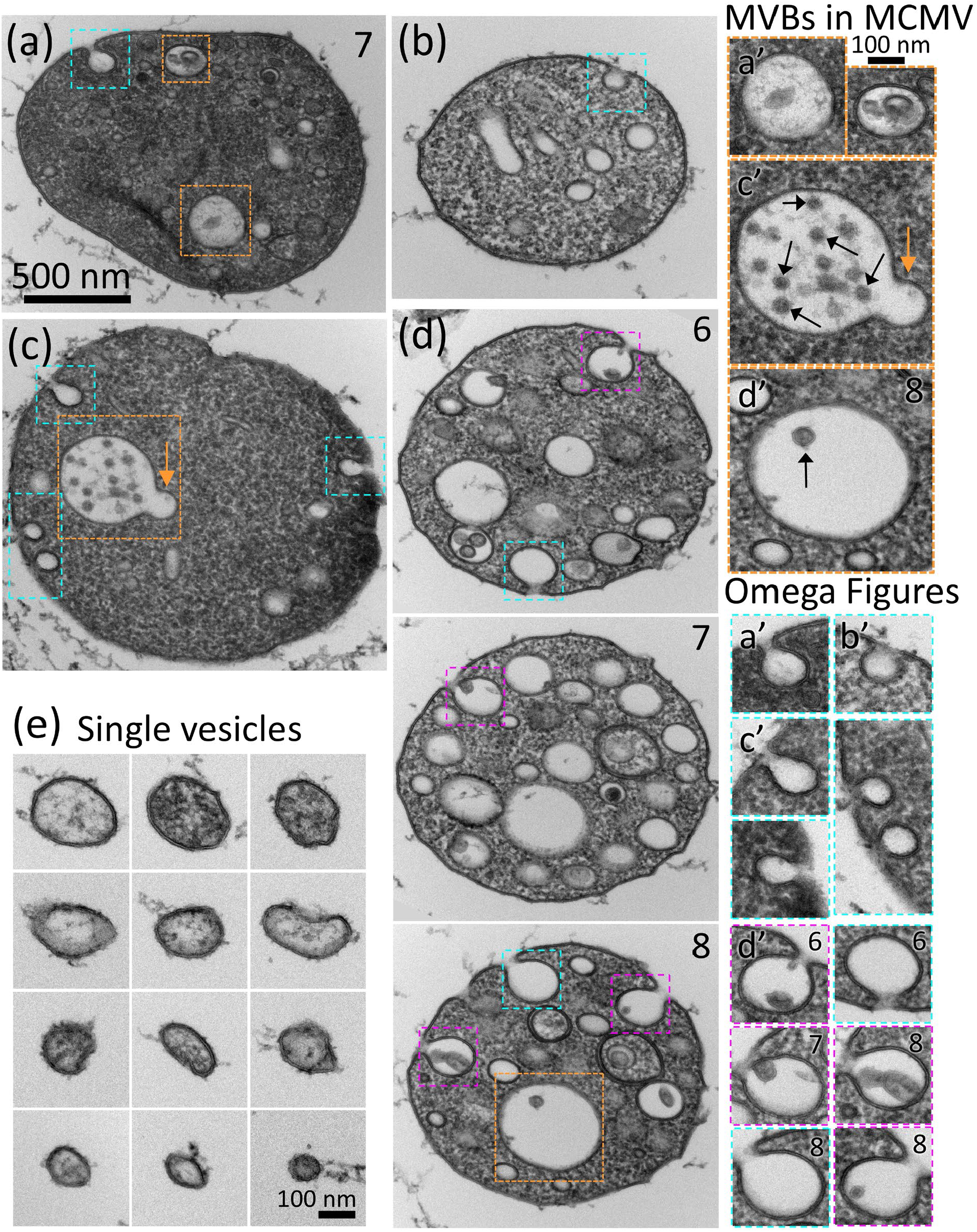
MCMVs contain MVBs that undergo remodeling and exocytosis of ILVs to release exosomes. (a, c, d) MCMVs containing organelles that are structurally identical to MVBs (orange boxes in a, c, and d section 8). The MVB boxed orange in (c) contains a several ILVs (upper c’, black arrows) and appears to undergo an outward budding remodeling step (orange arrow). Three serial sections are shown through the MCMV shown in (d). Section 8 contains an MVB with one ILV (upper d’, black arrow). (a-c) MCMVs with one or more omega-figures on their limiting membrane that do not contain an ILVs, indicating exocytic activity (turquoise boxes, enlarged in lower a’-c’). (d) Three serial sections of an MCMV in which exocytic events occur in each section, some of which contain ILVs (magenta boxes). The ILV inside the omega figure often appears stuck to the inner membrane of the exocytic profile (lower d, sections 6, 7, and 8). (e) Single-membrane vesicles located on the coverslip surface that are structurally like the inner vesicles of MCMVs. Scalebar in (a) applies to (a-d). Supplemental movies 6 and 7 show aligned sections through the MCMVs shown in (a) and (d), respectively.

The most stringent structural test for membrane fusion between organelles and the plasma membrane is the so-called ‘omega figure’, describing the shape of a fusion intermediate in a cross-section thinner than the fusion pore. Thus, if MCMVs secrete exosomes, omega figures of fusing MVBs should occasionally be visible on the MCMV limiting membrane, especially since the cultures were preserved by ultrafast freezing which occurs in milliseconds and can capture dynamic events like exocytosis.

Further examination revealed MCMVs with one or more omega-shaped profile on their limiting membrane (Figure 3 a-d and lower a’-d’). The profiles resembled smaller versions of the exocytic profiles of MVBs caught in the act of fusing with the cell plasmalemma in reticulocytes that also preserved by ultra-fast freezing and processed by freeze substitution and thin section EM (Harding, Heuser, and Stahl 1983).

If exosomes are released from MCMVs via these omega profiles, ILV-like vesicles should be seen inside or nearby the fusion event. Indeed, some of the omega figures contained ∼ 50-100 nm vesicles with grey lumens that were indistinguishable from ILVs (Figure 3d and lower d’, magenta boxes). Omega figures of fusion can span two sections and may appear empty in one section but seen to contain an ILV in the neighboring section (Supplemental figures 4 and 5, and Supplemental movies 6 and 7).

The ILVs still inside the omega figures appeared stuck to the inner membrane (Figure 3d and lower d’, magenta boxes) which apparently prevented their diffusion out. Interestingly, this observation made sense of the peripheral ILVs seen stuck to the outer surface of the MCMV in Figure 2d. It is likely that peripheral vesicles reveal a history of ILV exocytosis from the MCMV, and ILVs stuck on the inner MVB surface become exposed on the outside of the MCMV upon complete flattening of the omega figure. These observations are consistent with the idea that MVBs inside MCMVs fuse and release exosomes from MCMVs.

### 3.6 Single-membrane vesicles corresponding to MCMV’s internal vesicles are found on coverslip surface

If MCMVs release some of their internal vesicles, including ILVs (exosomes), they too might attach to the coverslip surface, and it might be possible to find single-membrane vesicles that match the size and appearance of MCMV inner vesicles in the first one or two sections cut along the surface of the coverslip. Such areas were searched at high magnification, and single-membrane vesicles ranging in size from ∼50-200 nm were found (Figure 3e). Like the vesicles observed inside MCMVs, the vesicles had grey smooth or granular interiors. The observation of these vesicles was consistent with their release from MCMVs, but they likely also arose from EV release directly from cells.

## 4 Discussion

Here a new class of microvesicle is described, termed MCMV, that buds from cellular protrusions clustered on the plasma membrane of HUVECs (Illustrated in Figure 4). The cellular membrane protrusions contain vesicular cargo that when compared with cytoplasmic organelles, could be identified as MVBs with ILVs, endosomes (round, tubular and clathrin coated), ER, and mitochondria. Serial sections showed that the protrusions are a few hundred nanometers to 1 micron thick and in some cases extended up in the Z axis relative to the coverslips for more than ∼1200 nm, beyond the scope of the serial sections analyzed. Protrusions were often branched and intertwined. At branch points and connections to the cell, the protrusions often became constricted to thin necks of 65-200 nm. Exploration of the coverslip surface between cells revealed MCMVs that were immobilized on the coverslip and contained MVBs containing ILVs in addition to other vesicle types. Preservation by fast-freezing captured omega figures formed by MVB-like organelles fusing with the MCMV limiting membrane, suggesting that some MCMVs secrete small vesicles that are structurally identical to ILVs. The omega figures on the MCMV-limiting membrane do not exhibit a clathrin coat, so are unlikely to be endocytic. It also seems of low probability, but not impossible, that ILV-sized vesicles are captured and internalized into and MCMV from the relatively vast volume of culture medium. Taken together, these observations suggest a novel pathway by which a subset of exosomes are released from a transiting MCMV after pinching off from a protrusion on the HUVEC surface.

**FIGURE 4:**
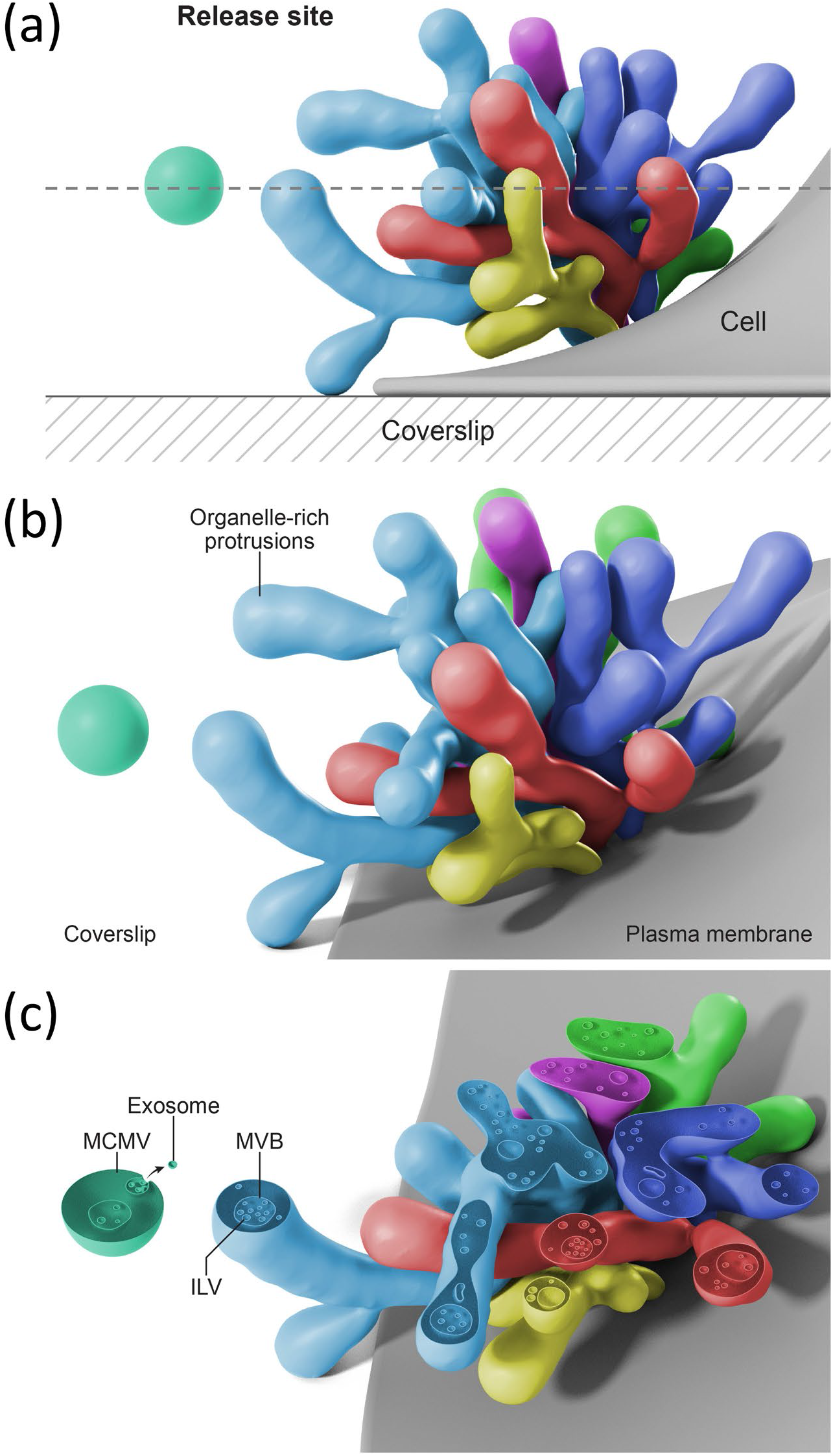
Representation of an MCMV release site from protrusions on a cultured endothelial cell and exosome secretion from a released MCMV. (a) Depiction of several protrusions (colorized in different colors) clustered on a cultured endothelial cell surface, and a nearby pinched off MCMV floating in the extracellular milieu. A slice plane in the *en face* orientation relative to the coverslip is represented by the dashed line. (b) A view at a slightly different angle shows that the protrusions are often branching and intertwined. (c) A slice through the protrusions and the MCMV shows internal round and tubular vesicular organelles, including MVBs containing ILVs, inside the protrusions and the MCMV. An omega figure on the MCMV limiting membrane indicates exosome secretion from the MCMV.

A previous scanning and transmission EM study of HUVECs described what is likely the same site of protrusions as we document and reported that vesicles shed from this specialized site on the plasma membrane contain proteases that promote angiogenesis (Taraboletti et al. 2002). Similarly, a separate scanning EM study showed discrete areas of membrane protrusions on otherwise relatively smooth surface membranes of unstimulated HUVECs (Combes et al. 1999). Indeed, various specialized domains of membrane protrusions have been shown to shed MVs on a number of cell types (Rilla 2021).

The rapid freezing, freeze substitution and serial section EM methodology utilized here (Walther and Ziegler 2002) is currently being applied to other cell types to determine if MCMVs are released by other cell types, or are unique to HUVECs. A growing number of cryo-EM studies document EVs containing internal vesicles derived from diverse sources such as human plasma, cerebrospinal fluid, urine, and ejaculate, a human mast cell line, and rodent neuronal culture media suggesting that release of MCMVs is a mechanism conserved in other cell types (Arraud et al. 2014; Brisson et al. 2017; Hoog and Lotvall 2015; Zabeo et al. 2017; Gamez-Valero et al. 2019; Emelyanov et al. 2020; Matthies et al. 2020). Similarly, EVs containing vesicles have been documented in some thin section EM studies of chemically fixed cells (Valcz et al. 2019).

Other cellular processes produce EVs containing internal vesicle compartments, similar to MCMVs. For example, cells undergoing apoptosis fragment into apoptotic bodies containing micronuclei and other organelles, and range in size from ∼500 nm to several microns in diameter (Xu, Lai, and Hua 2019). However, none of the MCMVs described in the current paper appeared to contain micronuclei. When undergoing apoptosis, adherent cells (including apoptotic HUVECs) show cell shrinkage, condensation of chromosomal DNA, fragmentation of nuclei, substrate detachment, and plasma membrane blebbing, which were rarely observed in our cultures.

A second class of EV that contains internal compartments is exophers, large (∼4 μm diameter) membrane-bound organelles which are extruded from touch neurons in *Caenorhabditis elegans* (Melentijevic et al. 2017) and murine cardiomyocytes (Nicolas-Avila et al. 2020) to expel potentially toxic materials such as dysfunctional mitochondria and protein aggregates. Additionally, in *C. elegans*, exophers released by muscle cells in pregnant females transfer nourishing yolk proteins to support developing embryos (Turek et al. 2021). Exophers are the largest class of EV (up to 15 μm), and characteristically contain numerous mitochondria which is not a distinguishing feature of MCMVs (Turek et al. 2021). There can also be a tube connecting the exopher to the cell from which it was extruded, but no connections were observed between MCMVs and cells.

Migrasomes are a third type of EV that contains numerous internal vesicles, and they perhaps share the most similarities with MCMVs, however migrasomes were shown not to contain MVBs (Ma et al. 2015). Migrasomes form by a migration-dependent mechanism at the termini and branch points of retraction fibers emanating from the trailing edge of migrating cells such as NRK cells (Ma et al. 2015) and other cell types and tissues (Di Daniele, Antonucci, and Campello 2022). However, MCMVs were not attached to retraction fibers, nor did they have “tails” of broken fibers as seen in electron micrographs of migrasomes (Ma et al. 2015; Yu and Yu 2021; Jiao et al. 2021). Also, remnants of retraction fibers were not observed on the coverslip in abundance, and the distribution of MCMVs on the coverslip surface appeared random relative to neighboring cells, and not concentrated on one side of a cell (potentially the trailing edge). Since the protrusions on HUVECs extend up from the cell in the Z-axis relative to the coverslip rather adhered to the coverslip substrate like retraction fibers, our interpretation is that MCMVs pinch off and diffuse in the culture medium before occasionally landing and becoming immobilized on the coverslip surface. It will be interesting to confirm this interpretation in future studies and determine if the location of the protrusion site on HUVECs correlates with the trajectory of cell movement.

Two aspects of MCMVs set them apart from these other classes of multi-compartmented EVs. First, MCMVs contain internal organelles that are identical to MVBs and ILVs observed in the cell cytoplasm. Second, exocytosis of ILVs from the internal MVBs was observed, thereby effecting exosome secretion from the MCMV. The direct observation of exocytosis of ILVs suggests a functionality that is unique to MCMVs (or has yet to be discovered in other types of EVs). It also suggests that the function of MCMVs is not removal of cellular debris, although this too could be a function as has been proposed for exophers and migrasomes. To be functional as signaling entities, EVs must deliver messages, in the form of bioactive molecules to recipient cells. Packaging cargos inside multiple layers of membrane rather than a unilamellar carrier could shield EV contents from degradation in the extracellular space, enabling them to voyage farther before being released from the MCMV or taken up into recipient cells. Multiple layers of membrane could also help vesicle contents avoid lysosomal degradation in the recipient cytoplasm and/or reach the nucleus. Additionally, grouping multiple vesicles of related signaling molecules in a unified EV could deliver contents as a component kit, rather than relying on coincidental arrival of components in separate EVs, at the right place and in the right quantities allowing for more efficient signaling.

Of note, in this study the HUVECs analyzed were at about 70% confluence to better observe potential sites of EV biogenesis on the cell peripheries. This raises the possibility that release of MCMVs occur in response to a “wound like”, sub-confluent state and perhaps participate in wound healing. Future studies are needed to determine if MCMVs are released from confluent HUVECs and from other types of endothelial cells in culture and *in vivo*.

A challenge in the study of EVs has been the isolation of EV subpopulations. Though possessing different sites of origin, MVs and exosomes share overlapping size ranges, molecular compositions, and densities, rendering biochemical enrichment and characterization of EV subsets a challenge (Kowal et al. 2016). The findings of this paper suggest that part of the difficulty may arise from EVs that consist of exosomes inside of microvesicles. Their presence could go unrecognized or misinterpreted as apparent overlap in biophysical properties. Moreover, attempts to separate microvesicles from exosomes may prove futile when MCMVs are both.

In summary, we have described a domain of protrusions extending from the plasma membrane of resting HUVECs that contain membrane bound organelles including MVBs. MCMVs bud from the protrusions and contain vesicular compartments, including MVBs that can fuse with the MCMV limiting membrane and release exosomes. MCMVs can be evaluated as a new type of organelle-containing microvesicle, and a potential source of exosome release that occurs remotely from the parent cell, adding new considerations to when, where, and how EVs are assembled and released from the endothelium and potentially other cells and tissues.

## ACKNOWLEDGEMENTS

This work was funded by the intramural programs of the NICHD (DIR) and the NCI Center for Cancer Research. The authors thank Stéphane Mahé (ThermoFisher Scientific) for technical support with the Tecnai T20 microscope and Ethan Tyler (Medical Arts Branch, NIH) for the artistic rendering. We also thank Hang Waters for critical reading of the manuscript.

## AUTHOR CONTRIBUTIONS

Writing-Original Draft: JDP

Conceptualization: JDP, JZ,

EM Methodology: JDP, JZ, EM, DDR, SK

Project administration: JZ

Formal analysis: JDP

Investigation: JDP, EM

Visualization: JDP

Writing-review & edit: DDR, EM, SK, JZ

Resources: JZ, DDR

Supervision: JZ, DDR

Funding acquisition: JZ, DDR

