## Supplemental Figures for "Endothelial cells release microvesicles that harbor multivesicular bodies and secrete exosomes"

**SUPPORTING INFORMATION**

**SUPPLEMENTAL FIGURES:**

**SUPPLEMENTAL FIGURE 1**

**SUPPLEMENTAL FIGURE 2**

**SUPPLEMENTAL FIGURE 3**

**SUPPLEMENTAL FIGURE 4**

**SUPPLEMENTAL FIGURE 5**

**SUPPLEMENTAL MOVIES:**

**SUPPLEMENTAL MOVE 1**

**SUPPLEMENTAL MOVIE 2**

**SUPPLEMENTAL MOVIE 3**

**SUPPLEMENTAL MOVIE 4**

**SUPPLEMENTAL MOVIE 5**

**SUPPLEMENTAL MOVIE 6**

**SUPPLEMENTAL MOVIE 7**

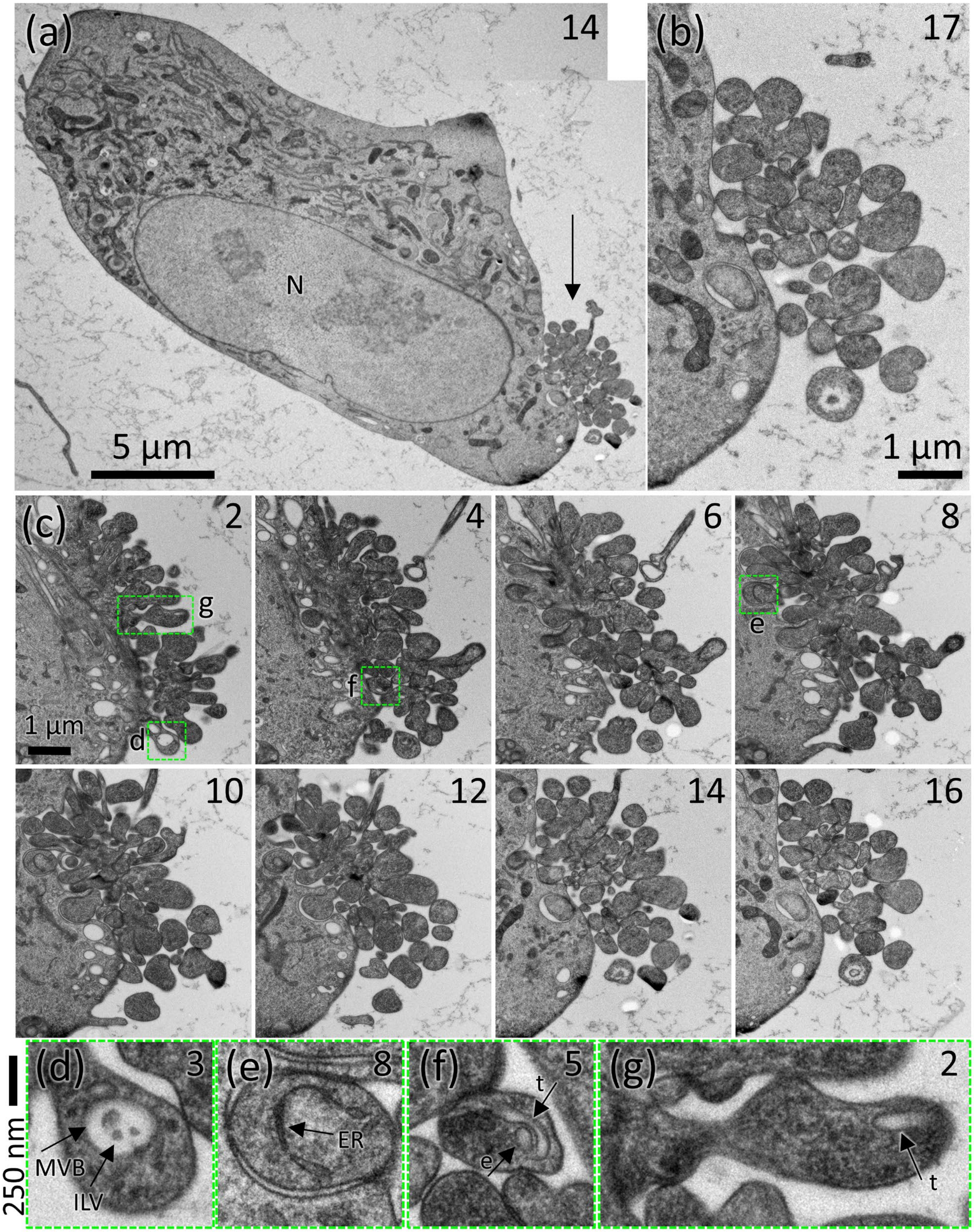
(a)
14
N
5 µm
(b)
17
1 µm
(c)
2
g
1 µm
d
4
f
6
8
e
10
12
14
16
250 nm
(d)
3
MVB
ILV
(e)
8
ER
(f)
5
t
e
(g)
2
t

**SUPPLEMENTAL FIGURE 1:** (a) A thin section through an entire HUVEC shows a smooth cell surface interrupted by site of protrusions localized to one end of the cell (arrow). Serial section number is indicated in the upper right corner of each image. The higher the number, the greater the distance from the coverslip. N, nucleus. (b) Enlarged view of the protrusion site as it appears in section 17, ~1200 nm above the surface of the coverslip. At this height, the protrusions are cut in cross-section and do not appear attached to the cell. (c) Even numbered sections through the protrusion site are shown with section number in the upper right corner. All sections can be viewed in Supplemental movie 2. Green boxed areas show protrusions that are enlarged in (d-g) with arrows indicating membrane-bound organelles including MVBs containing ILVs, endoplasmic reticulum (ER), round endosome (e), and tubular endosome (t).

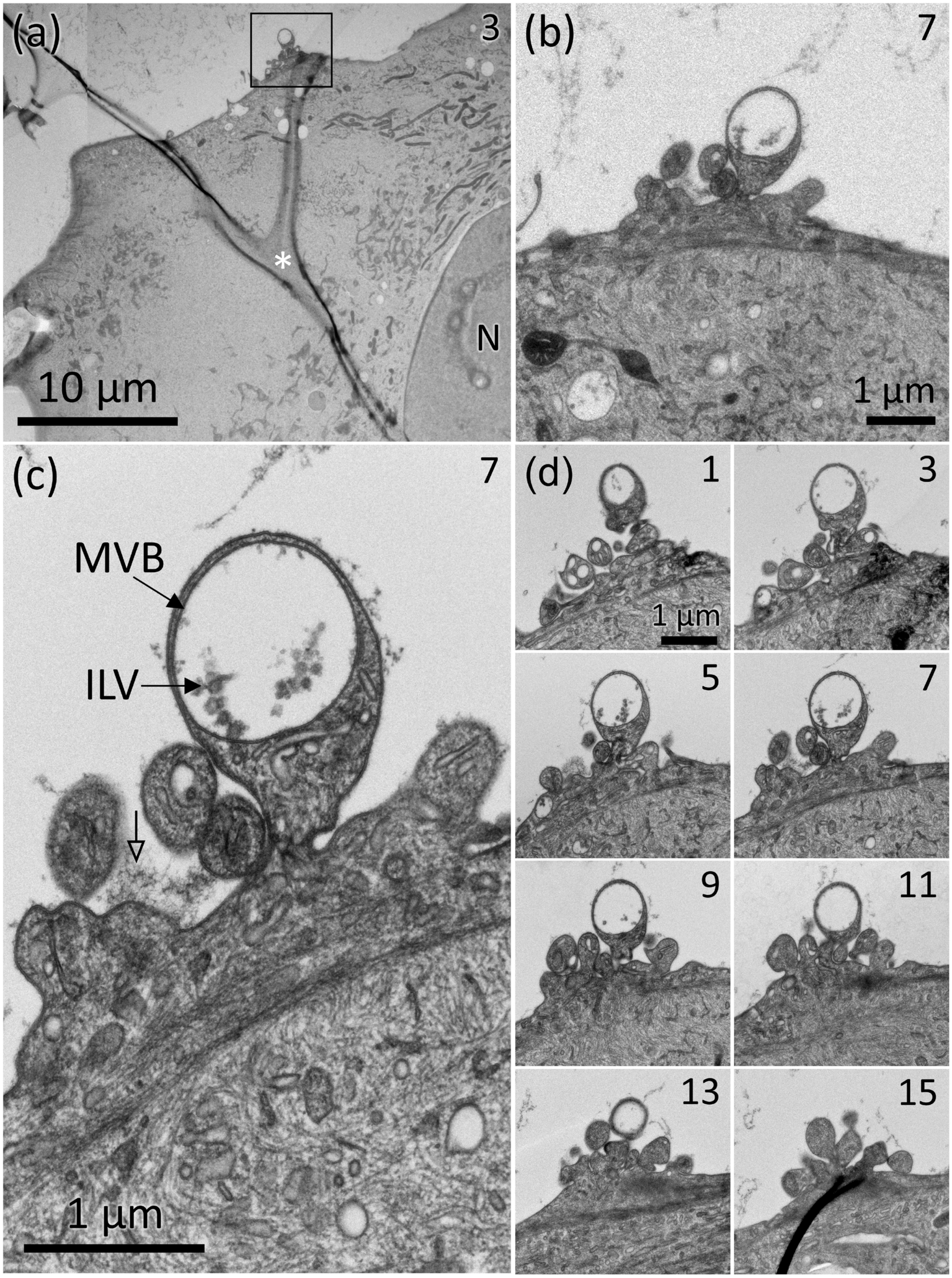
(a)
3
N
10 µm
(b)
7
1 µm
(c)
7
MVB
ILV
1 µm
(d)
1
1 µm
3
5
7
9
11
13
15

**SUPPLEMENTAL FIGURE 2:** (a) A site of protrusions (boxed) on the otherwise smooth surface of a HUVEC. White asterisk indicates a Y-shaped wrinkle in the section. N, nucleus. Serial section number is indicated in the upper right corner of each image. An enlarged view of the protrusion site boxed in (a) is shown in (b) and (c). Arrows indicate an MVB containing ILVs in a prominent bulb-shaped protrusion. Open arrowhead indicates fibrous material occasionally present between protrusions. (d) Odd-numbered serial sections through the protrusion site are shown, see also Supplemental Movie 3.

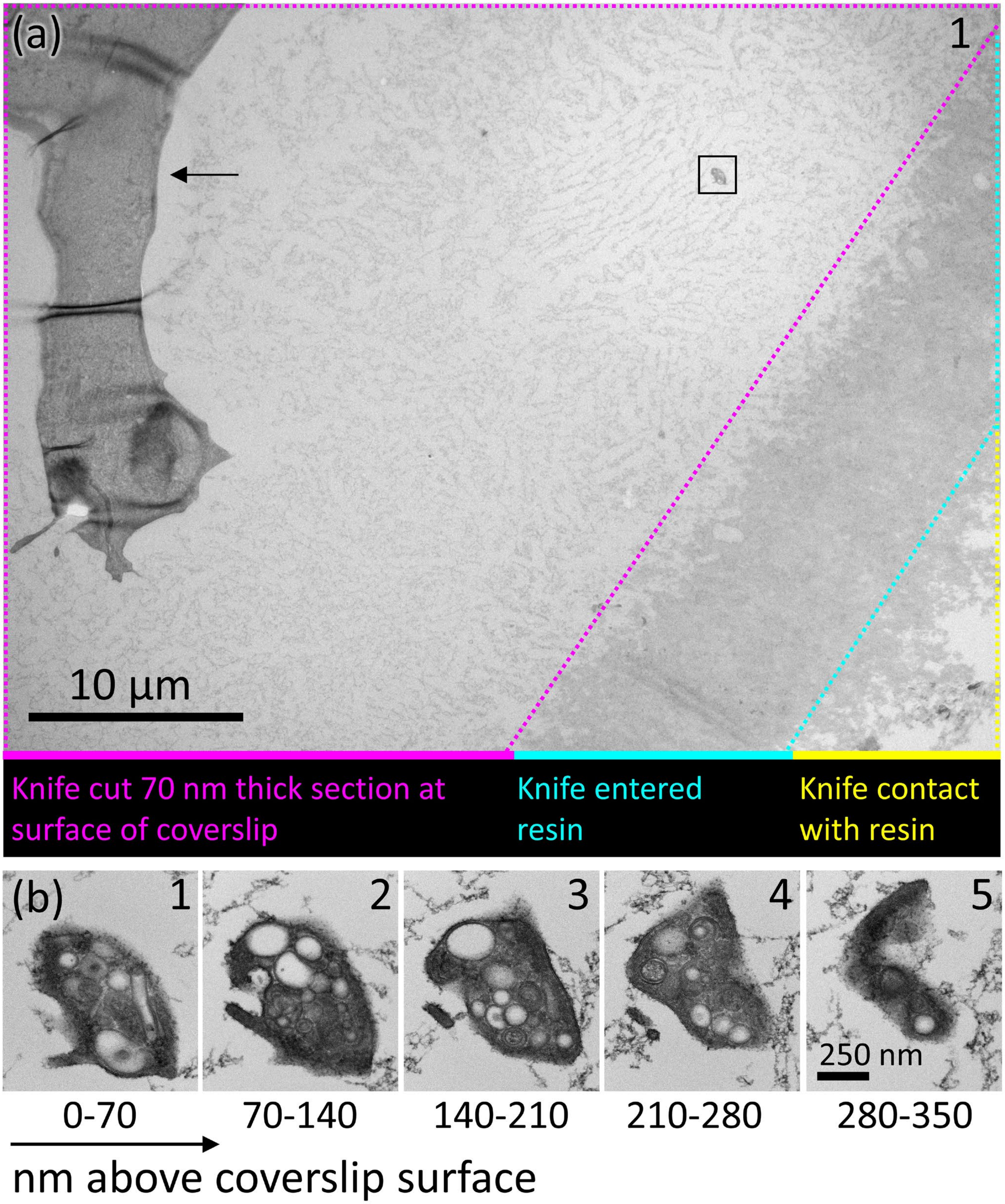
(a)
1
10 µm
Knife cut 70 nm thick section at surface of coverslip
Knife entered resin
Knife contact with resin
(b)
1
2
3
4
5
250 nm
0-70
70-140
140-210
210-280
280-350
nm above coverslip surface

**SUPPLEMENTAL FIGURE 3:** (a) Field of view showing an MCMV (boxed) located 25 µm from the nearest cell (arrow) and 10 µm from where the knife entered the resin during sectioning. The ragged edge (indicated by yellow boundary) is where the knife contacted the surface of the resin, then entered the resin (turquoise boundary), and then began cutting a 70 nm thick section (magenta boundary), confirming that the MCMV was located on the surface of the coverslip. (b) The MCMV is sectioned completely in five, 70 nm-thick serial sections, having a height of about 350 nm on the coverslip. See Supplemental movie 4 for higher magnification view of aligned images and the numerous vesicles and tubules inside.

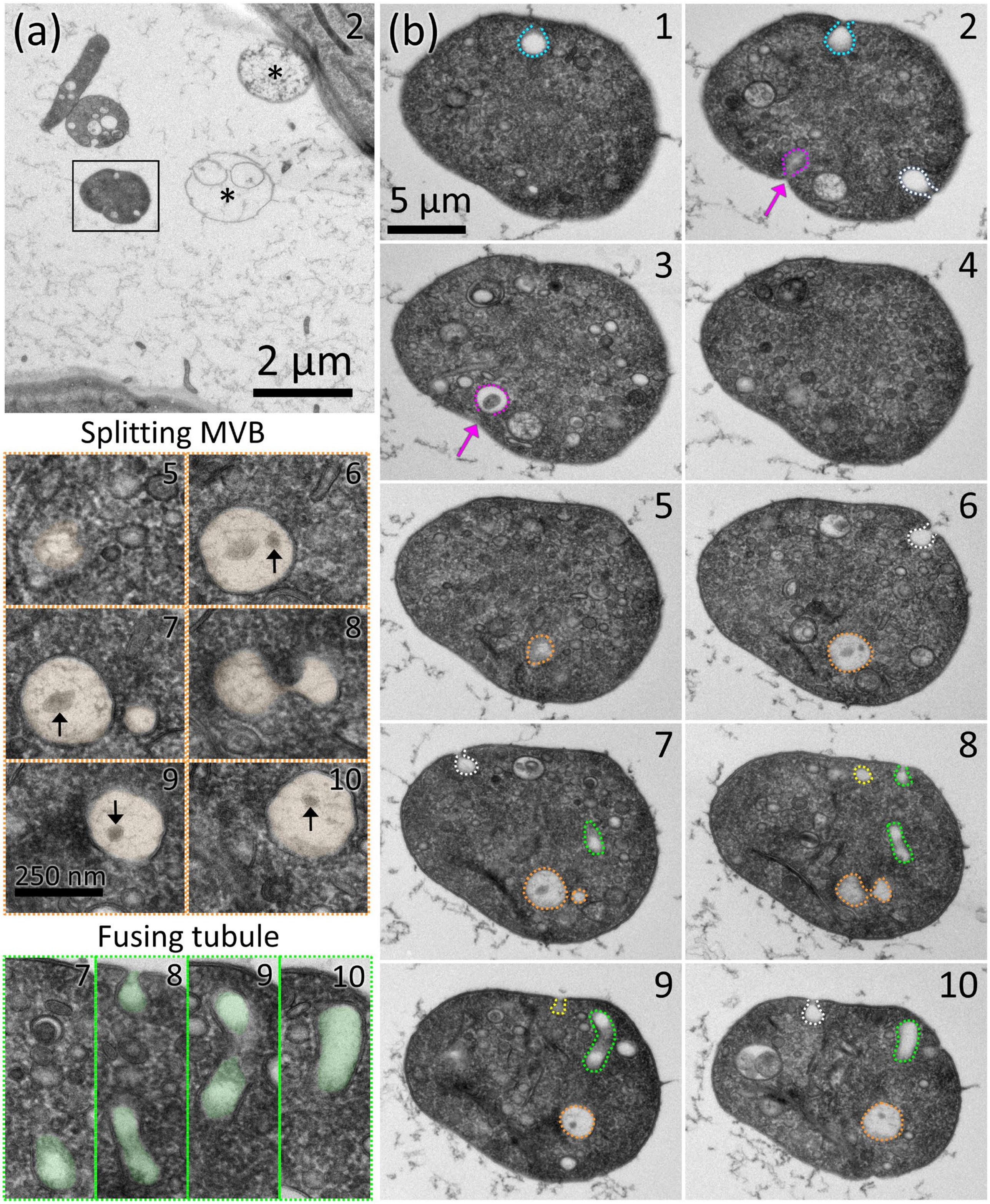

(a)
2
*
*
2 µm
Splitting MVB
5
6
7
8
9
10
250 nm
Fusing tubule
7
8
9
10
(b)
1
2
5 µm
3
4
5
6
7
8
9
10

**SUPPLEMENTAL FIGURE 4:** (a) Field of view showing the location on the coverslip of the MCMV in figure 3a (black box). Asterisks mark two structures that appear to be degraded MCMVs. Serial section number is indicated in the upper right corner of each image. (b) Ten sections through the MCMV boxed in (a). White dashed lines indicate omega figures that appear on one section. Turquoise and yellow dashed lines trace empty omega figures that span two sections. The magenta dashed lines trace an ILV-containing omega figure that spans two sections (magenta arrow indicates pore opening). Orange dashed lines outline an MVB-like organelle that contains four ILVs and appears to be splitting into two MVB-like organelles (shown enlarged to the left). Green dashed lines trace a curved tubule that goes in and out of sections 7-10, and in section 8 can be seen fusing with the MCMV periphery (shown enlarged to the left). Supplemental movie 6 shows 11 aligned sections through the MCMV shown in (b).

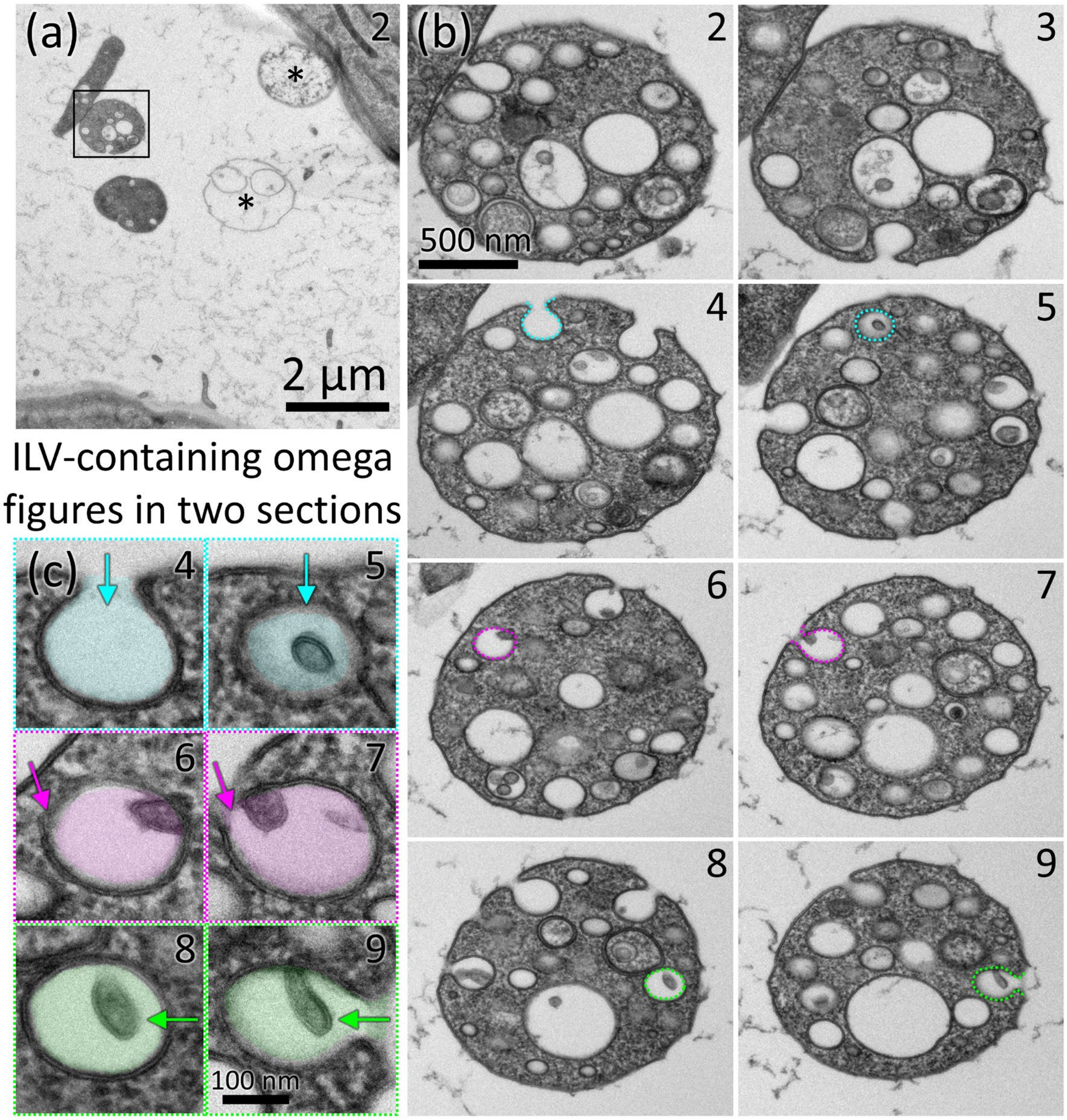


**SUPPLEMENTAL FIGURE 5:** (a) Field of view of the MCMV shown in Figure 3d (black box). The MCMV is near a tubule-shaped organelle that looks like an MCMV (not analyzed). Asterisks mark two structures that appear to be degraded MCMVs. Serial section number is indicated in the upper right corner. (b) Eight serial sections thought the MCMV. Colored dashed lines trace omega figures that span two consecutive sections demonstrating that an omega figure that appears empty in once section can contain an ILV (or second ILV) in the neighboring section. (c) Enlarged views of the color-coded omega figures shown in (b). Arrows indicate the position of the opening to the omega figure. Supplemental movie 7 shows 11 aligned sections through the MCMV shown in (b).

**SUPPLEMENTAL MOVIES**

**SUPPLEMENTAL MOVIE 1:** Aligned stack of serial sections of the protrusion site shown in Figure 1. Sixteen serial sections encompassing 1120 nm thickness in Z-height beginning at the surface of the coverslip. Note that section 9 is absent due to a fold in that section. Section number is indicated in the upper right corner of each image.

**SUPPLEMENTAL MOVIE 2:** Aligned stack of serial sections of the protrusion site shown in Supplemental figure 1. Seventeen sections encompassing 1190 nm thickness in Z-height. Section number is indicated in the upper right corner of each image.

**SUPPLEMENTAL MOVIE 3:** Aligned stack of serial sections of the protrusion site shown in Supplemental figure 2. Odd-numbered sections encompassing 1050 nm thickness in Z-height are shown. Section number is indicated in the upper right corner of each image.

**SUPPLEMENTAL MOVIE 4:** Aligned stack of five serial sections capturing the entire height of the MCMV shown in Supplemental figure 3 containing round and tubular internal vesicles. Section number is indicated in the upper right corner of each image.

**SUPPLEMENTAL MOVIE 5:** Aligned stack of ten serial sections capturing the entire height of the MCMV shown in Figure 2. Section number is indicated in the upper right corner of each image.

**SUPPLEMENTAL MOVIE 6:** Aligned stack of eleven serial sections through the MCMV shown in Figure 3a and Supplemental figure 4b. Many internal vesicles are smaller than the 70 nm thickness of the section, and thus appear in only one section. Section number is indicated in the upper right corner of each image.

**SUPPLEMENTAL MOVIE 7:** Aligned stack of eleven serial sections through the MCMV shown in Figure 3d and Supplemental figure 5b, showing numerous sites of exocytosis occurring on the MCMV periphery. Section number is indicated in the upper right corner of each image.
